# Reactivating Positive Personality Traits During Sleep Promotes Positive Self-Referential Processing

**DOI:** 10.1101/2022.11.27.518064

**Authors:** Ziqing Yao, Tao Xia, Jinwen Wei, Zhiguo Zhang, Xuanyi Lin, Dandan Zhang, Pengmin Qin, Yina Ma, Xiaoqing Hu

## Abstract

Positive self-view is evident by a bias in favor of positive self-referential processing, as individuals tend to endorse positive characteristics over negative ones when making self-judgments. While research suggests that a positivity bias can contribute to psychological well-being, it remains unclear how to enhance positive self-referential processing. Here, we reported an integrated training procedure that aimed at enhancing individuals’ positive self-referential processing. Specifically, participants engaged in a cue-approach training task (CAT) during wakefulness where they gave speeded motor responses to positive personality traits. In a subsequent nap, we unobtrusively re-played half of the trained positive traits during participants’ slow-wave sleep to reactivate memories associated with these positive traits (targeted memory reactivation, TMR). Upon awakening, we found that CAT+TMR enhanced participants’ positive self-referential processing, as evidenced by faster endorsement of positive traits. Further analysis revealed that this enhancement was associated with specific brainwave patterns during sleep: delta (1–4 Hz) traveling waves moving from posterior to anterior brain regions. These findings demonstrate the potential benefits of integrated wakeful cue-approach training and sleep-based memory reactivation in strengthening positive self-referential processing.

## Introduction

People often perceive themselves through rose-tinted lenses, exhibiting a positivity bias (Taylor & Brown, 1988; Zell et al., 2020). This positivity bias is evident in self-referential judgments, as people preferentially choose positive personality traits to describe themselves and have better memories for positive traits compared to negative ones (Taylor & Brown, 1988; Watson et al., 2007; Guenther & Alicke, 2010; Romero et al., 2016; Dainer-Best et al., 2017; Collins & Winer, 2023). This positive self-referential bias is commonly associated with lower levels of depressive symptoms (e.g., self-doubt, worthlessness), and is crucial for mental well-being, especially when facing self-threatening information (Sowislo et al., 2013). While the psychological benefits of positive self-referential processing is well-established (Taylor & Brown, 1988; Colombo et al., 2020; Orth et al., 2022; Weisenburger et al., 2023), a significant gap exists in understanding how to effectively enhance this process (Orth et al., 2022; Hoffmann et al., 2023). To address this gap, we integrated two procedures that may enhance positive self-referential processing: (1) wakeful cued-approach training (CAT, Schonberg et al., 2014), and (2) a sleep-based targeted memory reactivation procedure (TMR, Oudiette and Paller 2013).

The CAT task prompts participants to give speeded motor responses to cued stimuli, ultimately increasing positive evaluations or preference toward these trained stimuli (Schonberg et al., 2014; Salomon et al., 2018; Schonberg & Katz, 2020; Itzkovitch et al., 2022). While CAT has been used to alter preferences for various stimuli, such as food, abstract patterns, images (for a review, see Salomon et al. 2018), its impact on higher-order social-cognitive processes such as self-referential processing remains unexplored. Complementing the wakeful CAT, the TMR aims to promote memory consolidation during post-training sleep, a phase vital for stabilizing newly acquired memories. During sleep, covert, repeated memory reactivation contributes to memory consolidation, notably during non-rapid eye movement (NREM) sleep characterized by the <4 Hz slow-wave activity (Diekelmann & Born, 2010; Rasch & Born, 2013; Klinzing et al., 2019; MacDonald & Cote, 2021; Brodt et al., 2023). TMR entails replaying memory-related sensory cues to sleeping participants, further strengthening episodic memories or even changing subjective preferences during NREM sleep (Creery et al., 2015; Hu et al., 2015; Schreiner & Rasch, 2015; Cairney et al., 2016; Ai et al., 2018; Lewis & Bendor, 2019; Abdellahi et al., 2023; for a meta-analysis of TMR, see Hu et al., 2020). Here, in the context of self-referential processing, we hypothesize that the integration of wakeful CAT and sleep-based TMR could change how individuals perceive and endorse positive personality traits as self-descriptive.

Specifically, while CAT primes the brain to be more receptive to specific positive traits, TMR works to consolidate these traits during sleep, potentially leading to more robust and enduring positive self-referential judgments. Therefore, we tested the joint impact of CAT and TMR on the positive self-referential processing, particularly focusing on how they influence the immediate and long-term endorsements and retention of positive personality traits.

During NREM sleep, cardinal neural oscillations such as slow oscillations (<1 Hz), delta waves (1–4 Hz) and the 12-16 Hz spindles are instrumental in mediating memory reactivation and consolidation (Born & Wilhelm, 2012; Rasch & Born, 2013; Antony et al., 2017; Klinzing et al., 2019; Schreiner et al., 2021; Petzka et al., 2022; Brodt et al., 2023). Specifically, in TMR, researchers repeatedly found that the cue-elicited delta and sigma EEG power changes predicted TMR benefits (Lehmann et al., 2016; Blume et al., 2017; Göldi et al., 2019; Denis & Payne, 2023; Liu et al., 2023; Schechtman et al., 2023; Xia et al., 2023). While these findings have significantly advanced our understanding, they remained silent on how propagation of sleep EEG oscillations may contribute to memory consolidation. The propagation of EEG oscillations across different brain regions, known as traveling waves, has been increasingly recognized for their significance in linking brain function to behavior (Muller et al., 2018). Importantly, slow oscillations and spindles have been observed to manifest as robust traveling waves propagating across the cortices (Muller et al., 2018). Such traveling waves may coordinate cross-region information flow and neural communications during sleep, which can be crucial for reactivating and consolidating memory traces (Massimini et al., 2004; Murphy et al., 2009; Hangya et al., 2011; Kurth et al., 2017). However, to date, how delta slow-wave activity as traveling waves may modulate memory is not yet understood. Given that the TMR employs external sensory cues to trigger internal memory reactivation, it is plausible that upon processing auditory cues such as the spoken positive traits, the delta slow waves would exhibit a forward trajectory from the posterior to the anterior frontal brain regions, contributing to effective processing of positive traits and thus influencing positive self-referential processing. Indeed, recent research suggests that a forward patterns of traveling wave could aid in the bottom-up processing of stimuli in a wakeful state (Alamia et al., 2023). We thus hypothesize that during TMR, the forward slow traveling wave activity could align sensory processing with semantic processing of positive traits during sleep, thereby enhancing positive self-referential processing.

Here, we employed an adapted version of the well-established self-referential encoding task (SRET) to quantify participants’ self-referential processing (Derry & Kuiper, 1981; Dainer-Best et al., 2017, 2018; Collins & Winer, 2023, for procedure and tasks, see Figure 1). In addition to this SRET, we assessed participants’ recall of self-referential traits from the SRET in a free recall task, and self-referential preferences in a probe task. To examine the immediate and possible long-term effects of TMR, we measured participants’ self-referential processing twice: immediately after the TMR and one-week later. Our findings revealed that the integration of CAT and TMR facilitated the endorsement speed of positive personality traits immediately after sleep. Moreover, analysis of cue-elicited EEG showed that the strength of 1–4 Hz forward traveling waves predicted the endorsement speed of positive traits during immediate test and the endorsement of positive traits one-week later.

**Figure 1.**
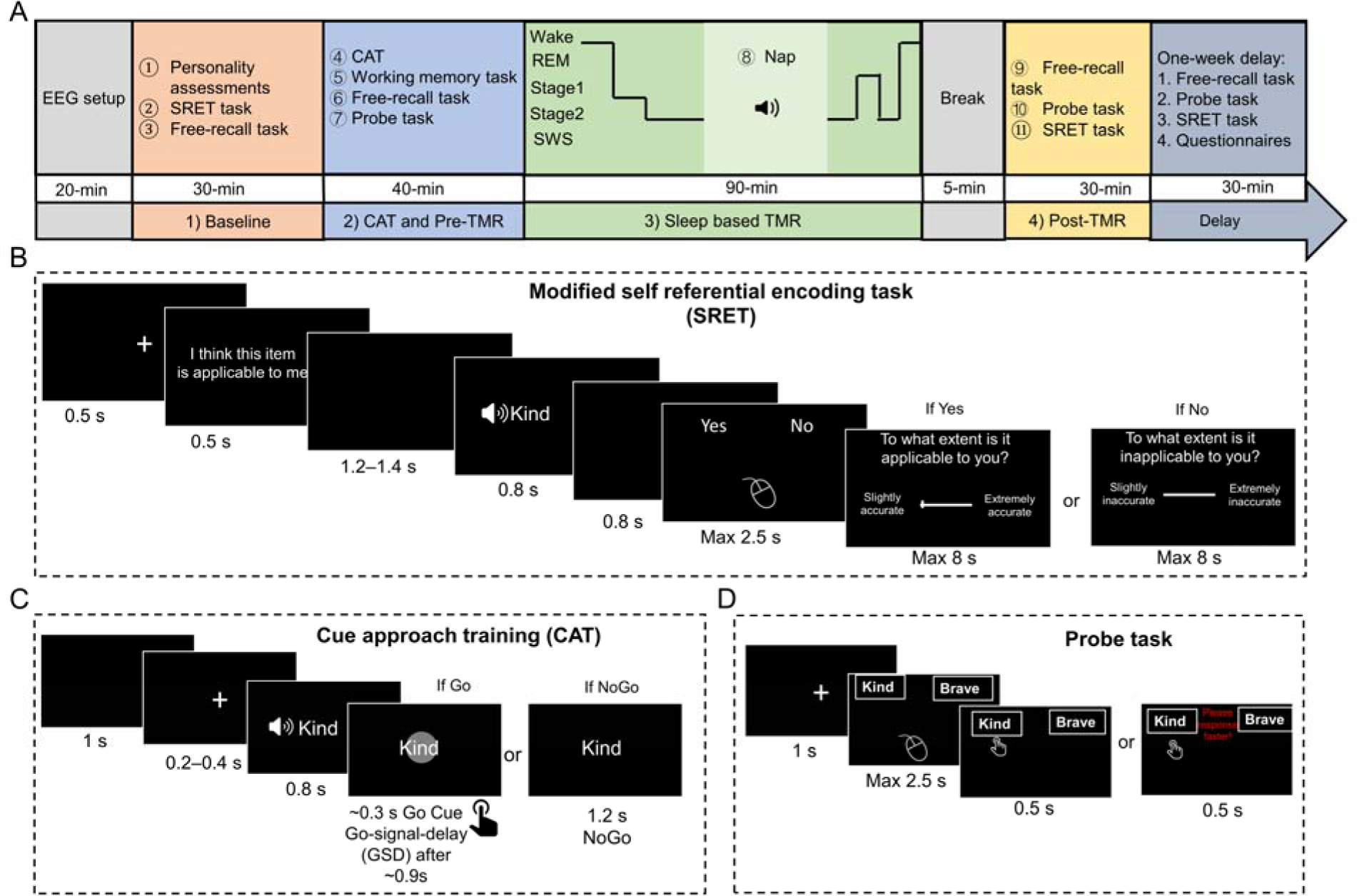
An overview of experimental design and main tasks. (A) The task flow illustrates the baseline tests (phase 1), CAT and post-CAT tests (phase 2), sleep-based TMR (phase 3), and post-TMR tests (phase 4), followed by a delayed tests phase after one week (*n* = 35). (B) Modified SRET, in which participants made speeded binary endorsement task to determine whether a personality trait was descriptive of oneself, followed by rating the accuracy of specific traits in describing themselves within the same trial (i.e., endorsement level). After completing the baseline SRET, participants performed a self-referential free recall task. In both the post-TMR and the one-week delay phases, participants completed the free recall task and the SRET with binary endorsements while omitting ratings. (C) An exemplar trial of CAT, in which participants either passively viewed positive traits presented visually and aurally (i.e., NoGo trials) or pressed a button when they saw a white circle appear immediately after the positive trait onset (i.e., Go trials). The average GSD (go-signal-delay) was 900 ms. (D) Probe test, participants were presented with pairs of positive Go and NoGo traits and were asked to select which trait was more self-descriptive. Note that Go and NoGo traits in each pair were matched on baseline self-descriptive ratings (see Methods for a full description of the procedure and experimental tasks).

## Results

### Awake CAT promoted self-referential preferences

First, to examine whether CAT promoted the preferences of positive Go traits, we analyzed the proportion of trials in which participants preferred Go traits over NoGo traits as better self-descriptive in the probe task (Figure 1D), using a generalized linear mixed model with participant factor as a random effect (GLMM, see Method for specific model). In each Go/NoGo pair, both traits had comparable initial endorsement level based on the baseline SRET rating phase. Consistent with previous CAT research (Salomon et al., 2018), we found that participants were more likely to choose Go over NoGo traits despite their comparable baseline endorsement level: mean proportionLJ=LJ53.3% (vs. chance level 50%), odds ratio (OR)LJ=LJ1.24, 95% CI [1.09, 1.41], *p* = 0.001. This result suggested that the CAT specifically increased participants’ self-referential choice of the Go traits in the probe task.

### Awake CAT + sleep TMR enhanced positive self-referential endorsement and speed

Having established the effectiveness of CAT in enhancing the preferences of positive Go traits, we next examined how TMR may further influence positive self-referential processing. Specifically, we analyzed two outcome variables from the SRET task, including binary endorsement choice, and reaction times (RTs) when endorsing positive traits. Note that we applied false discovery rate (FDR) corrections for all multiple comparisons.

To examine positive self-referential endorsements change, we ran a GLMM using baseline endorsement rating value as a covariate, TMR condition (Go-cued, Go-uncued, and NoGo-uncued) and time (baseline, post-TMR, and delay) as fixed effects, and participant factor as a random effect to predict endorsement choices (yes, no) of positive traits. Our results revealed a significant TMR effect, L (2) = 8.02, *p* = 0.018. Particularly, participants endorsed more Go-cued traits than NoGo traits (*p* = 0.026). No differences were found between other conditions (all *p*s >0.12). However, neither time nor the TMR by time interaction were significant, L (2) = 2.52, *p* = 0.284, L (4) = 2.80, *p* = 0.592, respectively.

Given that choice speed could indicate preferences (Konovalov & Krajbich, 2019), we analyzed item-level RTs when participants endorsed positive traits via an LMM including TMR conditions and time as fixed effects. The results showed a significant main effect of time, F (2, 35) = 8.37, *p* = 0.001, indicating that RTs became faster from baseline to post-TMR and delay (all *p*s < 0.001), while no difference was found between post-TMR and delay (*p* = 0.126). No significant main effects were observed for the TMR condition, F (2, 4921) = 0.73, *p* = 0.480. Notably, we found a significant TMR by time interaction, F (4, 4922) = 3.19, *p* = 0.013 (Figure 2A). Post-hoc comparisons revealed that in the post-TMR, participants were significantly faster in endorsing Go-cued traits than NoGo-uncued traits (*p* = 0.022). In contrast, this pattern was not observed in the delay testing (*p* = 0.414). Other comparisons across different time points did not yield any significant differences in RTs (Go-uncued versus NoGo-uncued, Go-cued versus Go-uncued traits, all *p*s >0.1). Taken together, these results suggest that the CAT+TMR jointly facilitated endorsement speed for Go-cued positive traits compared to NoGo-uncued traits.

**Figure 2.**
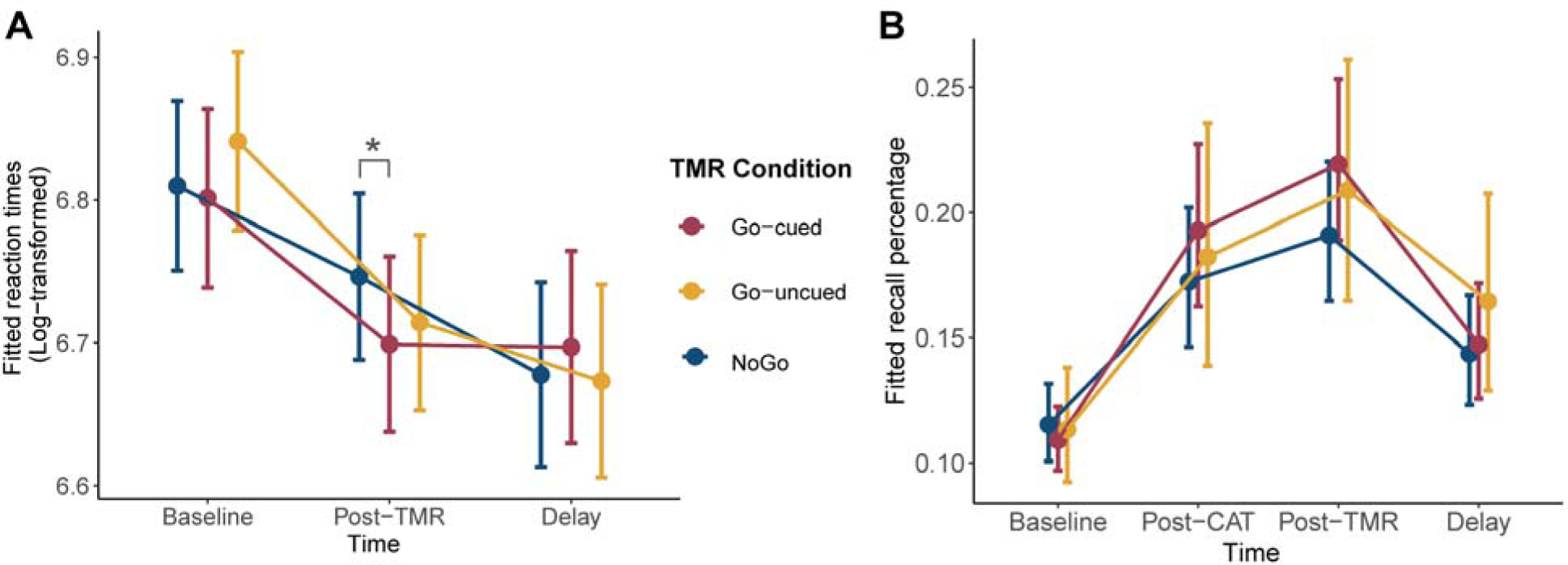
Behavioral results across time in the SRET tasks. (A). Fitted values for the interaction effect of TMR conditions and time on predicting log-transformed RTs for endorsing positive traits during the SRET. Error bars indicate 95% confidence intervals (CIs). (B). Fitted values for the interaction effect of TMR conditions and time on predicting recall percentage for positive traits during the SRET. * = *p* < 0.05.

To address the concern that the observed RT differences might be solely due to the influence of CAT, we conducted an additional LMM in two additional independent behavioral samples to investigate if RT differences in endorsing positive traits were present between Go and NoGo trait words. These samples consisted of one group undergoing active CAT with ‘Go’ training trials and another group exposed to passive CAT without such active components (See SOM for behavioral sample details). In this analysis, we included CAT conditions (Go vs. NoGo), time (baseline, post-CAT, and delay), and group (active vs. passive) as fixed effects, with participants as random effects, focusing on positive endorsement RTs. The results only showed a significant main effect of time, F (2, 75) = 12.62, *p* < 0.001, with faster endorsements after CAT and during the delay tests compared to the baseline (all *p*s < 0.001). However, we observed no significant main effects or interactions specifically attributable to CAT (all *p*s >0.35). These findings, therefore, suggest that it is the combination of CAT and TMR, rather than CAT alone or solely repetition of positive trait words, that promotes the endorsement speed of positive traits.

### Overall enhancement of positive self-referential memory recall following CAT and TMR

Recognizing the established efficacy of TMR in enhancing memory performance (Hu et al., 2020), our study specifically investigated whether integrating CAT with TMR would improve the recall of positive traits in self-referential memory tests. We used a GLMM including TMR (Go-cued, Go-uncued, NoGo-uncued) and time (baseline, post-CAT, post-TMR, delay) as fixed effects, and participant factor as a random effect to predict percentage of positive traits participants recalled as self-referential. Results revealed a significant time effect, L (3) = 72.70, *p* < 0.001, with post-hoc comparisons indicating that compared to baseline, there were significantly higher recalls of positive traits at post-CAT, post-TMR and delay tests (*p*s <0.001, Figure 2B). Moreover, recall declined from the post-CAT to the delay (*p* = 0.011) and from the post-TMR to the delay (*p* < 0.001), while there was no difference between post-CAT and post-TMR recalls (*p* = 0.063). Furthermore, we found no significant TMR effect, χ² (2) = 1.04, *p* = 0.595, nor any significant TMR by time interaction, χ² (6) = 2.29, *p* = 0.891, indicating that CAT or TMR did not selectively change recall performance. Lastly, when examining the recall of negative traits, the time effect was not significant, χ² (3) = 3.35, *p* = 0.341, suggesting that the increased recall was specific to positive traits.

To ascertain whether CAT+TMR jointly contributed to the overall enhancement of positive self-referential memory, we also contrasted the current CAT+TMR group with the abovementioned two additional independent behavioral samples (active CAT and passive CAT groups, see SOM for behavioral sample details). A subsequent GLMM, accounting for earlier recall as a covariate, showed a significant group effect, χ² (2) = 9.29, *p* = 0.010. Participants who received both TMR and CAT exhibited superior recall of positive traits compared to the passive CAT group (*p* = 0.008), but not higher than the active CAT group (*p* = 0.085), while no significant difference was found between the active and passive CAT groups (*p* = 0.229, all FDR corrected). Together, these result highlights that combining CAT and TMR had long-lasting impact in facilitating self-referential recall of positive traits.

Together, the findings demonstrated that CAT shifted preferences towards positive traits, while combining CAT and subsequent TMR effectively enhanced positive self-referential processing by accelerating RTs when for endorsing positive traits.

### Auditory processing of positive traits during sleep TMR

To first validate the processing of spoken positive traits during sleep, we quantified cue-elicited event-related potentials (ERPs) and time-frequency resolved EEG power changes during the TMR. Consistent with prior TMR research (Schreiner et al., 2015; Lehmann et al., 2016; Antony et al., 2018; Schechtman et al., 2021; Abdellahi et al., 2023; Guttesen et al., 2023; Liu et al., 2023; Schechtman et al., 2023; Xia et al., 2023), cue-elicited ERPs showed two positive clusters over frontal-central electrodes (F1/2, FC1/2, C1/C2, Fz, Cz) from 0.29 to 0.52 seconds and from 1.04 to 1.40 seconds (two-tailed *t*-test, cluster-based permutation-corrected *p* < 0.005). In addition, the time-frequency analysis also identified two significant positive clusters over frontal-central electrodes: the delta–theta–alpha band (1 to 12 Hz, 0 to 1.66 seconds), and the sigma– beta band (10 to 30 Hz, 0.3 to 1.42 seconds, two-tailed *t*-test, cluster-based permutation-corrected *p* < 0.001, Figure 3A, B). We next examined whether EEG power changes within these clusters may be associated with changes in RTs and choices during the positive self-referential processing. However, we did not find significant associations (all *p*_corrected_ > 0.562 for delta-theta-alpha; and > 0.364 for sigma-beta.

**Figure 3.**
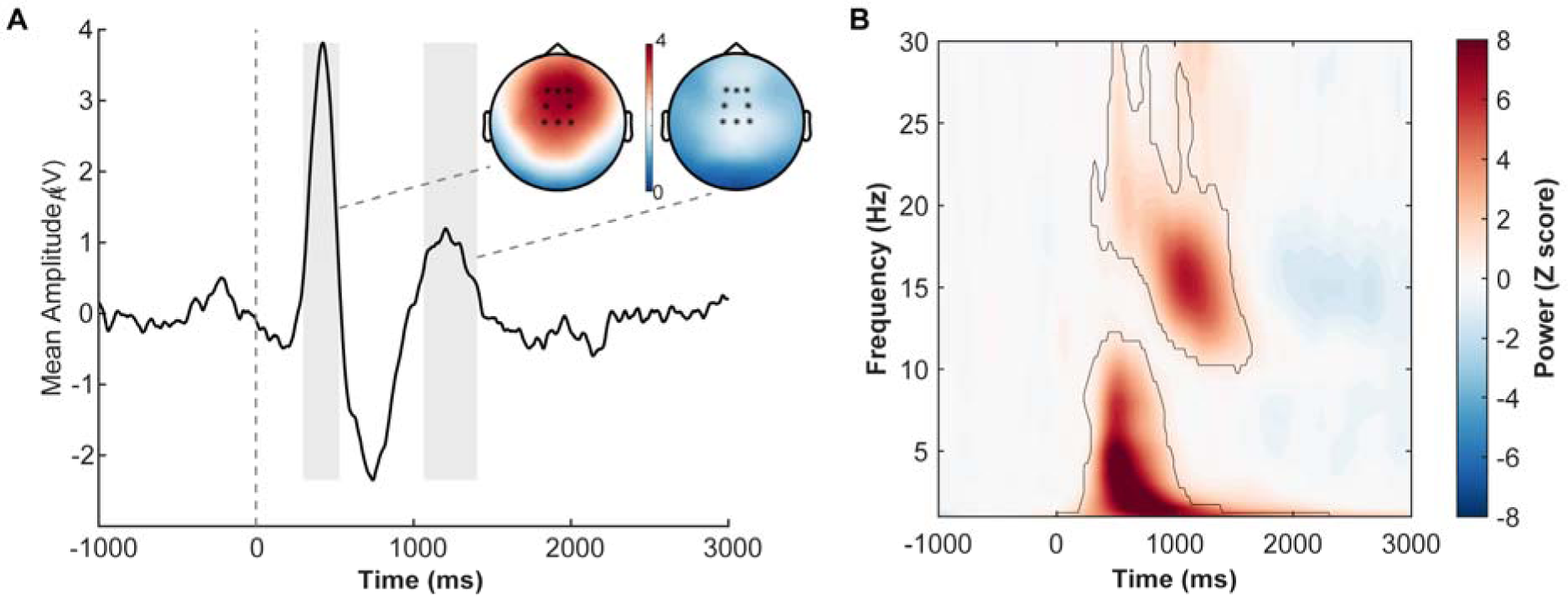
Cue-elicited power changes did not predict post-TMR endorsements or positive endorsement speed. (A) Grand averaged ERPs across frontal-central electrodes (F1/2, FC1/2, C1/2, Fz, Cz). Shaded area indicates significant time point when comparing ERPs against zero. Top right panel presents group average scalp topography of ERPs in response to TMR cues; with black circles highlighting the electrodes used in the ERP analysis. (B) Contour plot depicting the temporal and spectral characteristics of the significant clusters. Cluster a represents the low-frequency delta-theta-alpha band (1–12 Hz), and cluster b represents the sigma-beta band (10–30 Hz), with both clusters showing significant changes across the TMR time course (cluster-based permutation-corrected *p* < 0.001).

### Cue-elicited slow travelling waves predicted post-TMR positive traits endorsement speed

Next, we investigated how traveling waves during TMR might influence post-TMR positive self-referential processing. Given that TMR benefits would begin with effective sensory processing (posterior cortex), followed by high-level memory processing (see Liu et al. 2023), we hypothesized that the forward traveling waves propagating from posterior to anterior cortex would be critical here. Building upon methodologies used in previous research (Alamia et al., 2023), we analyzed the strength of directionality in both forward (from posterior to anterior brain regions) and backward (from anterior to posterior brain regions) traveling waves within the first 2s following the TMR cue onset, using EEG signals from midline electrodes (POz, Pz, CPz, Cz, Fz, FPz) during SWS in the 1–4 Hz frequency band (Figure 4A-C, see Method). The strength of forward and backward traveling waves was then utilized to predict item-level binary endorsement choices and RTs for endorsing positive traits, across immediate and a one-week delay test time, with False Discovery Rate (FDR) corrections for multiple comparisons.

**Figure 4.**
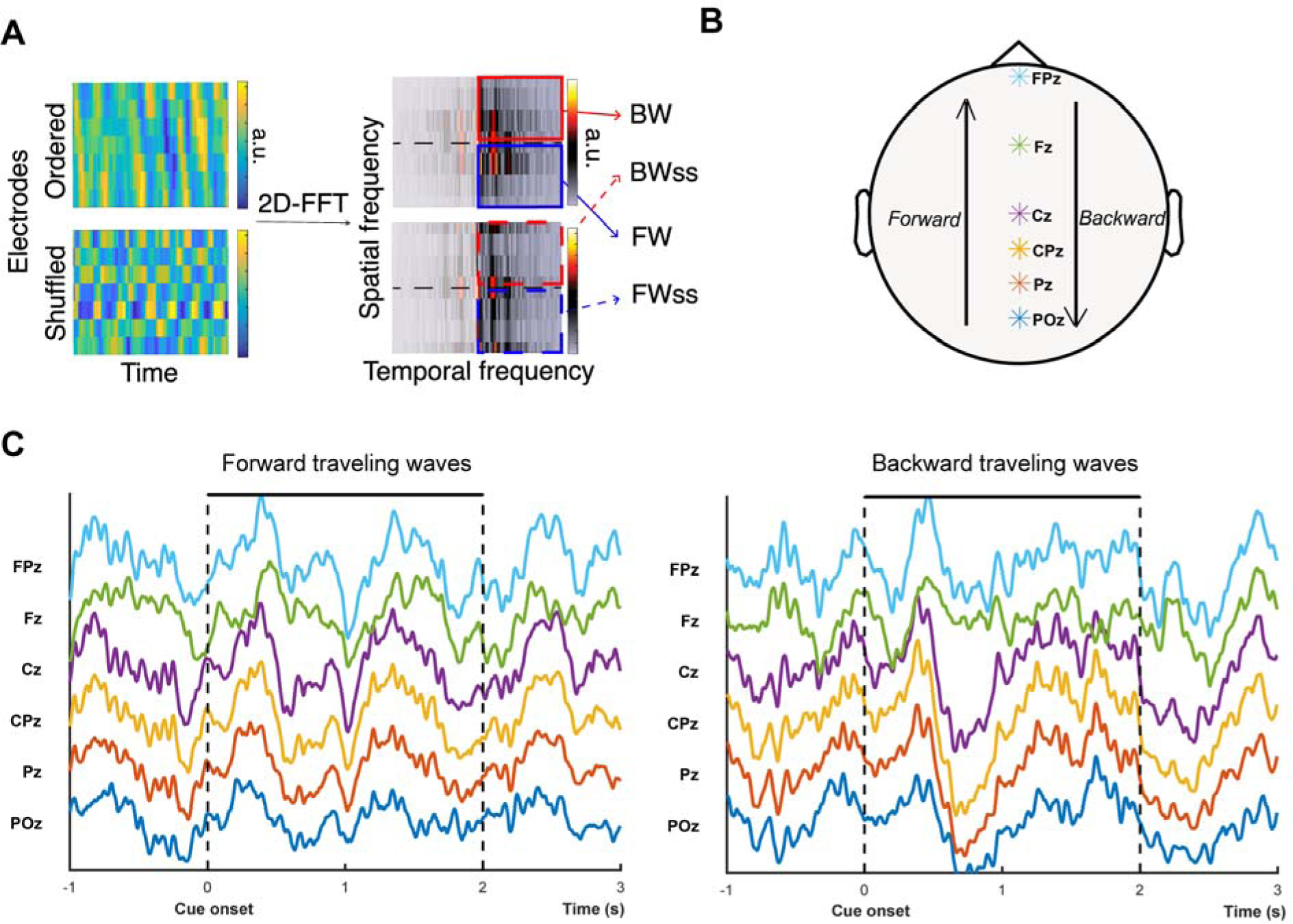
Slow travelling waves after TMR cue onset. (A). EEG signals from midline electrodes (0–2000 ms post-TMR cue) underwent two-dimensional Fourier transform (2D-FFT) decomposition, yielding power spectra. Baseline-corrected spectra using shuffled data delineate backward (anterior to posterior) and forward (posterior to anterior) traveling waves (detailed methodology in Alamia et al (2023). (B). Scalp diagram illustrating slow wave directionality. Forward waves are indicated by arrows pointing to frontal electrodes, backward waves to posterior electrodes. Asterisks mark analyzed electrode locations. (C) Demonstrations of forward (left panel) and backward (right panel) traveling waves.

We used (G)LMM with intensities of traveling waves as a fixed effect, alongside the number of trait repetitions during TMR and baseline endorsement ratings as covariates, and participant as a random effect, to predict post-TMR endorsements and RTs when endorsing positive traits. Results revealed that during the post-TMR test, neither forward nor backward traveling waves significantly predicted endorsement probabilities (*p*_corrected_ > 0.272, Figure 5A&B). Notably, forward traveling waves negatively predicted RTs when endorsing positive trats in the immediate test (*p*_corrected_ = 0.002, Figure 5C), but not in the one-week delay (*p*_corrected_ = 0.766, Figure 5G). Backward traveling waves showed no significant associations (*p*_corrected_ = 0.686, Figure 5D&H). Interestingly, forward traveling waves also significantly predicted the endorsement of positive traits after one week (*p*_corrected_ = 0.026, Figure 5E), suggesting a delayed effect on positive self-referential processing. Backward waves, however, did not demonstrate a significant prediction (*p*_corrected_ = 0.060, Figure 5F). Together, our results showed that the forward traveling waves elicited by positive traits during sleep can predict post-sleep positive self-referential processing.

**Figure 5.**
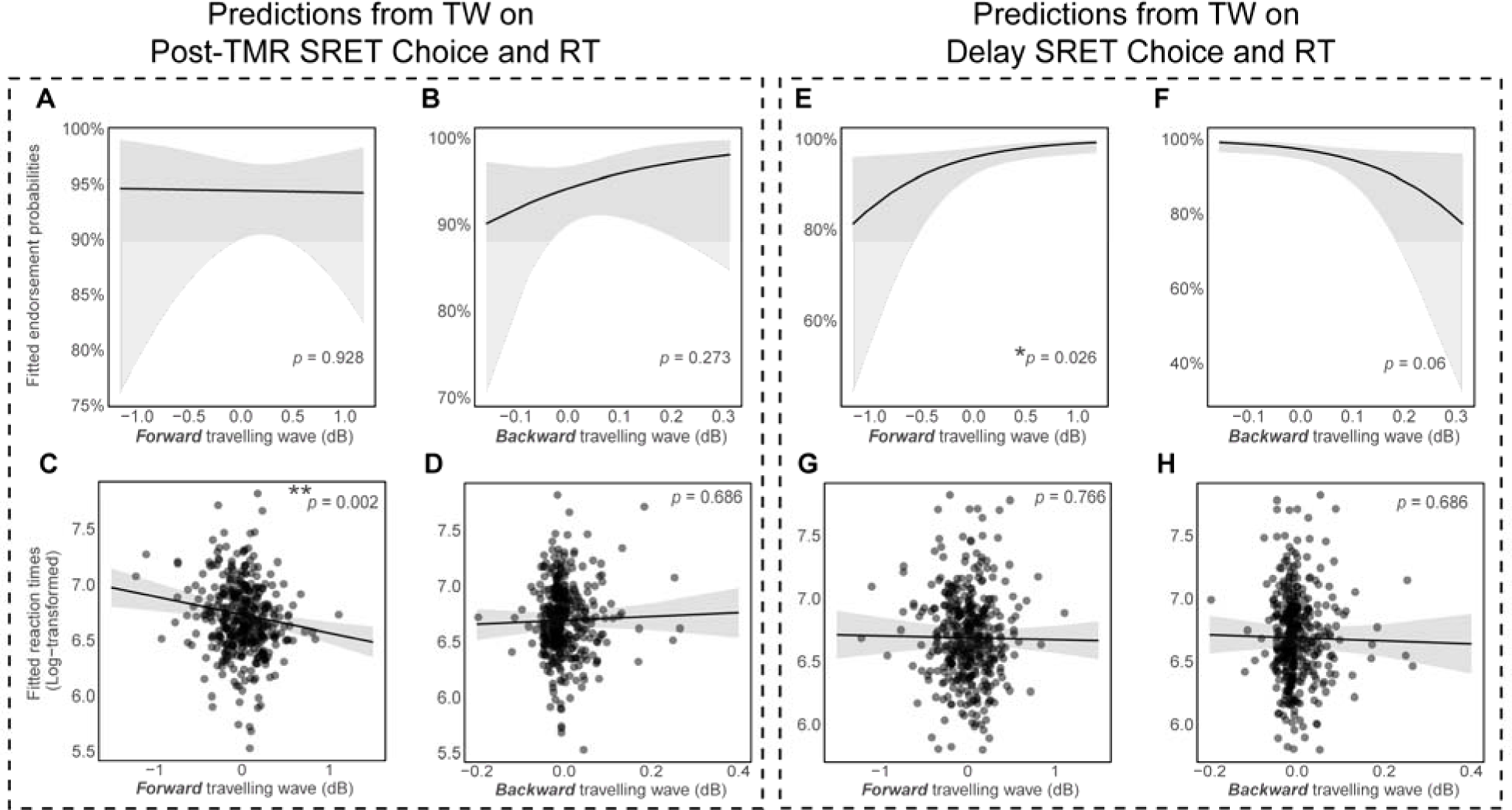
Predictive impact of delta traveling waves on endorsement probabilities and RTs in post-TMR and delay phases of the SRET task. Predictions from forward and backward traveling waves on (A, B) endorsement probabilities in post-TMR phase. (C, D) RTs in post-TMR phase. (E, F) endorsement probabilities in delay phase. (G, H) RTs in delay phase. In panel (C, D, G, H), each data point corresponds to the fitted value from a single trial within the LMM. Where data points overlap, they present a darker shade. In panel (A, B, E, F), we excluded raw data points due to their binary (zero or one) nature. Shaded area indicates 95% confidence intervals (CIs). TW: traveling wave. SRET: self-referential encoding task. ** = *p* < 0.01. * = *p* < 0.05.

## Discussion

By combining wakeful cue-approach training (CAT) and sleep-based targeted memory reactivation (TMR), we found that this integrated procedure effectively enhanced participants’ positive self-referential processing. We first used CAT to heighten participants’ preferences for specific “Go” positive traits, extending the existing CAT research. Following this, TMR was employed to re-play a subset of these Go traits during participants’ SWS sleep, further enhancing their accessibility and consequently promoting positive self-referential processing. The CAT and TMR expedited endorsement of these Go-cued positive traits, although it did not selectively alter self-referential memory immediately after sleep TMR. After one week delay, we observed a general increase in positive memory recall, rather than TMR-specific changes. Additionally, the presence of 1–4 Hz forward slow travelling waves during TMR was associated with enhanced positive self-referential processing, indicating an important role of cross-regional forward neural communications in driving behavioral benefits. These new findings contributed to our understanding of how to modulate and enhance positive self-referential processing.

We first found that the CAT successfully increased participants’ likelihood to choose Go over NoGo traits as self-descriptive in the probe task, demonstrating CAT’s efficacy in influencing self-referential choices. This finding extends the known effects of CAT on consumables such as snacks (Salomon et al., 2018; Schonberg & Katz, 2020), revealing its capability to shape high-level self-referential processing. Following the CAT phase, we replayed a subset of the trained positive traits during sleep to determine the cumulative impact of CAT and TMR on self-referential processing. Behaviorally, we found that participants exhibited faster endorsement of positive traits when these traits were trained during the wakeful CAT and subsequently reactivated during the SWS (Go-cued), compared to untrained traits (NoGo-uncued). This contrast highlights the joint benefits of CAT and TMR in facilitating the speed of positive self-referential processing. Apart from positive endorsement speed, we also observed that participants endorsed more Go-cued positive traits than NoGo-uncued traits as self-descriptive independent of testing time. Previous CAT research has indicated that the CAT can improve stimulus salience, effectively making these traits more prominent during the waking state (Schonberg et al., 2014; Schonberg & Katz, 2020). TMR during post-training sleep, on the other hand, further promoted memory reactivation and consolidation via cueing, improving the accessibility and retrieval efficiency of the cued stimuli (Walker & Stickgold, 2006; Diekelmann & Born, 2010; Klinzing et al., 2019; Lewis & Bendor, 2019; Brodt et al., 2023). In the context of our study, the TMR following the CAT likely further augments the accessibility of the trained traits, thereby speeding up their endorsements.

The accelerated endorsement of positive traits and the overall higher endorsement probabilities for CAT+TMR traits can be partially explained by neural oscillations during sleep and TMR. In contrast to power spectral analysis that often concentrates on regional oscillations, the concept of traveling waves encompasses a wider array of neural characteristics. These include both spatial propagation and frequency property, offering a more comprehensive view of the spatial-temporal dynamics of brain activity during sleep (Massimini et al., 2004; Muller et al., 2018; Zhang et al., 2018; Halgren et al., 2019). Our study observed that the 1–4 Hz delta forward traveling waves, moving from posterior to anterior brain regions within 2 seconds after cue onset, were significantly associated with post-TMR endorsement RT and the endorsement of positive traits following a one-week delay. This finding, for the first time, suggests that delta forward traveling waves are pivotal in memory consolidation during sleep, broadening the existing knowledge of the neural mechanisms supporting sleep-mediated memory consolidation (Massimini et al., 2004). Previous studies have indicated the thalamic origin of delta waves during sleep (Adamantidis et al., 2019) and its integral role in memory consolidation (Schreiner et al., 2022). Our results contribute to this body of knowledge by demonstrating that the delta waves, as traveling waves, may also contribute to memory consolidation during exogenous memory reactivation. Furthermore, the forward direction of delta traveling waves implies the bottom-up processing of external cues in TMR sleep. Specifically, re-playing spoken positive traits during sleep, may initiate basic auditory processing at the posterior brain regions that advanced to high-level semantic and self-referential processing at the frontal regions. This forward propagation of delta slow waves may effectively integrate the positive traits into one’s self-scheme, contributing to positive self-referential processing. Although the exact neurocomputing processing remains elusive, parallels can be drawn from awake-state studies. A noteworthy study highlighted the bottom-up processing of visual stimulus is facilitated by alpha-band forward traveling waves (Alamia et al., 2023). These waves spatiotemporally organize distributed brain areas, enabling efficient processing of external stimuli. Consequently, we proposed that the sleeping brain processes spoken positive traits by coordinating different brain regions through traveling waves, particularly the forward delta traveling waves that propagated from posterior to anterior brain regions.

When evaluating participants’ self-referential memories using a free recall task, we did not find significant main effects of CAT or TMR. This result may stem from the experimental design, where participants engaged in the free recall task twice prior to sleep. Repeated recall may induce fast memory consolidation that makes the self-referential memories less susceptible to TMR (Antony et al., 2017; Liu et al., 2023). Notably, a week later, participants showed an overall enhanced memories for positive traits compared to the baseline, regardless of CAT or TMR manipulations. Supporting this, broader sleep and TMR studies indicate memory enhancement can often be observed over extended periods (Rakowska et al., 2021; Barner et al., 2023). Even more intriguingly, we found that this non-selective, general memory enhancement was only observed in the CAT+TMR group, but not in the other two groups (CAT only, or passive CAT). These findings suggest that the post-CAT sleep and TMR may further enhance the overall positive self-referential memories, regardless of cueing. Indeed, previous TMR research suggested that memory reactivation during sleep may have generalized benefits: in addition to enhancing cue-specific memories, TMR also strengthened uncued memories that shared the same context as the cued memories, leading to overall benefits of both cued and uncued memories (Schechtman et al. 2023; see also Oudiette et al., 2013 for TMR generalization effects).

Future directions and limitations shall be discussed. First, our study follows most prior research in administering the TMR during the NREM sleep, given the established link between NREM sleep and TMR benefits (see Lewis and Bendor 2019; Hu et al. 2020). However, research also pinpoints the role of REM sleep in modulating emotional memory and vocabulary learning (Batterink et al., 2017; Hutchison et al., 2021). Future research could investigate how TMR during REM sleep, and how the REM-related neural activity may impact the consolidation of self-referential memories. Second, while positive self-referential processing is linked with mental wellness (Wisco, 2009; Lou et al., 2019; Collins & Winer, 2023), our study did not examine how our procedure may impact outcomes that bear direct clinical relevance, such as depression-related symptoms. Future research is warranted to investigate whether enhancing positive self-referential processing may directly alleviate depressive symptoms (Orth et al., 2022; Hobbs et al., 2023). Third, while our research question concerns self-referential memory, we did not include non-personal traits as a control condition. Future studies could consider including a control task, specifically designed to disentangle non-self-referential memory from self-referential memory in understanding the CAT and TMR effects.

In conclusion, our study presents a novel approach in enhancing positive self-referential processing by combining wakeful motor training and sleep-based memory reactivation. In addition to behavioral benefits, our findings underscore the importance of cue-related forward delta traveling waves in predicting the speed of endorsing positive traits, establishing a direct connection between traveling waves induced by TMR and positive self-referential processing. By reinforcing positive self-referential processing through CAT+TMR, it may be possible to alter maladaptive cognitive biases or restore self-esteem, contributing to improved mental health outcomes.

## Supporting information

SOM

## Acknowledgements

The research was supported by the Ministry of Science and Technology of China STI2030-Major Projects (No. 2022ZD0214100), National Natural Science Foundation of China (No. 32171056), General Research Fund (No. 17614922) of Hong Kong Research Grants Council, Key Realm R&D Program of Guangzhou (No. 20200703005) to X.H.

## Author contributions

Ziqing Yao: Conceptualization, Data curation, Formal analysis, Investigation, Methodology, Project administration, Writing – original draft, Writing – review & editing. Tao Xia: Formal analysis, Visualization, Writing – review & editing. Jinwen Wei: Formal analysis, Writing – review & editing. Zhiguo Zhang: Writing – review & editing. Xuanyi Lin: Writing – review & editing. Dandan Zhang: Writing – review & editing. Pengmin Qin: Writing – review & editing. Yina Ma: Funding acquisition, Writing – original draft, Writing – review & editing. Xiaoqing Hu: Conceptualization, Methodology, Funding acquisition, Resources, Supervision, Writing – original draft, Writing – review & editing.

## Data and code availability

The data and analytical code supporting the study’s findings are available at the Open Science Framework repository: https://shorturl.at/bEG23.

## Experimental model and study participant details

### Participants

Our final sample included 35 participants with valid behavioral and EEG data (8 males, Mage ± SD = 20.83 ± 2.20 years), which is comparable to recent TMR studies (e.g., Schechtman et al. 2023). Nine additional participants had inadequate number of cues (< =3 rounds) due to relatively short slow-wave sleep (SWS). To ensure signal-to-noise ratio in the EEG analyses, experiments for these participants were terminated following the TMR, and data from these nine participants were not included in subsequent analyses. An additional participant was excluded because he or she reported hearing the cues during sleep. To facilitate sleep in the lab, we asked participants to wake up one hour earlier than their usual waking time and to avoid consuming caffeinated drinks on the day prior to – and of – the experiment. Participants were pre-screened regarding any current or history of sleep, psychiatric, or neurological disorders and had normal or corrected-to-normal vision. Participants received monetary compensation for their time (250 RMB, ∼36 USD), and gave written consent prior to the experiment. The study was approved by the Human Research Ethics Committee of the University of Hong Kong.

### Materials

All experimental procedures were implemented in E-Prime® 3.0 (Psychology Software Tools, Inc., Sharpsburg, Pennsylvania, USA). A pilot group of 20 participants rated personality traits (two characters trait words) on a scale from 1 (extremely negative) to 9 (extremely positive). We selected 60 positive personality trait adjectives (e.g., ‘clever’, M ± SD = 6.92 ± 0.44) and 60 negative personality trait adjectives (e.g., ‘lazy’, M ± SD = 3.00 ± 0.44; see SOM for the complete list of personality traits). Each spoken trait lasted around 1 second (range: 0.72-1.08s, M ± SD = 0.91 ± 0.08s). During the TMR phase, we used a neutral trait (valence rating: 4.9) as a control word.

### Method details

#### Task overview

Participants attended two lab sessions, scheduled approximately one week apart. In the first session, participants arrived to the lab at approximately 12:00 pm (exact arrival times ranged between 11:30 am to12:30 pm), where they read and signed consent forms and were set up with EEGs. Subsequently, a series of four task phases began in which participants completed a number of tests, beginning with baseline tests in the first phase, followed by CAT and post-CAT tests in the second phase, sleep-based TMR in the third phase, and post-TMR tests in the fourth phase. In the preliminary baseline phase, participants completed computer-based personality questionnaires, serving as a cover story for the personality trait words (hereafter, traits) presented to them in the following SRET. During the SRET, participants rated the extent to which specific traits described themselves. Participants then completed a self-referential free recall test. In the second phase, participants manually responded to positive traits (i.e., Go traits), prompted by visual and aural cues presented on screen and from a nearby loudspeaker (CAT). Participants then completed a free recall test and a probe test, in which they were presented with Go and NoGo trait word pairs and asked to select the trait word that was more self-descriptive. In the third phase, half of the positive traits were aurally re-played to sleeping participants during slow-wave sleep (SWS). Then, in the fourth phase, participants completed the same free recall test, probe test, and SRET. In the second lab visit (∼ 7 days later), participants completed the same free recall test, probe test and SRET as previously completed in the final phase of the first visit to examine the possible long-term TMR effects. Thus, they completed four self-referential free recall tests (baseline, post-CAT, post-TMR, delay), three SRETs (baseline, post-TMR, delay), and three probe tasks (post-CAT, post-TMR, delay).

#### Baseline tasks

Participants completed preliminary computer-based personality questionnaires, including the Rosenberg Self-Esteem Scale (RSES) (Rosenberg, 1965), Narcissistic Personality Inventory (NPI) (Raskin & Hall, 1981), Big Five Inventory (BFI) (John et al., 1991), Beck Depression Inventory-II (BDI-II) (Beck et al., 1996), State-Trait Anxiety Inventory (STAI state and STAI trait) (Spielberger, 1983), and Barratt Impulsiveness Scale (BIS-11) (Patton et al., 1995). Completing these questionnaires served as a cover story for the subsequent self-referential encoding task (SRET): participants were told that the personality traits that would be presented in the SRET were from their questionnaire data (for descriptives, see Table S1).

In the SRET (see Figure 1B), a cross symbol was presented on a computer screen at the beginning of each trial for 0.5 seconds, followed by the presentation of the sentence ‘I think this word is applicable to me’ in the center of the screen for another 0.5 seconds. After 1.2 to 1.4 seconds, participants were presented with a random word, given visually in written form and aurally from a speaker, from a selection of 120 adjectives for 0.8 seconds. After, participants were shown a blank screen for another 0.8 seconds and then were prompted to select if a trait word applied to them within 2.5 seconds by moving the mouse cursor continuously. The spatial location of ‘Yes’ and ‘No’ responses were counterbalanced (upper left/upper right or upper right/upper left). Following a ‘Yes’ response, participants were asked to rate the extent to which a trait word applied to them on a scale ranging from “slightly accurate” to “extremely accurate”, covertly equating to values from 1 to 50; following a ‘No’ response, participants were asked to rate the extent to which a trait word did not apply to them on a scale ranging from “slightly inaccurate” to “extremely inaccurate”, covertly equating to values from −50 to −1.

Following the SRET, participants completed a self-referential free recall task. Unlike previous free recall tasks wherein participants wrote down as many traits as possible, here, participants were asked to recall only the traits they had been presented with and they endorsed (i.e., “yes” response) during the previous SRET. Participants typed each recalled trait on a computer one at a time. Therefore, performance during this version of the recall task reflected self-referential memories.

#### Traits selection in the probe task

For each participant, we ranked all 60 positive traits in ascending order based on their baseline endorsement ratings (from 1, being the lowest rating and least self-descriptive, to 60, being the highest rating and most self-descriptive). We next equally divided these 60 traits into ‘Go’ and ‘NoGo’ trials, forming 30 Go-NoGo pairs for each participant. We chose traits for each pair based on each trait rating’s rank orders (i.e., from 1 to 60), to ensure that the Go and NoGo traits had comparable baseline ratings (*p* = 0.64, for details, see SOM and Figure S1A). For the post-TMR probe task, these Go/NoGo pairs were further categorized into cued (Go-cued) and uncued (Go-uncued) conditions, with each condition having 15 trait pairs. Full details for the trait allocations in the CAT and the probe task are provided in Supplementary Figure 1.

#### CAT and post-CAT tests

Following baseline assessments, participants completed a cue-approach training (CAT) task (see Figure 1C). For each CAT trial, a positive trait was presented visually and aurally for 1.2 seconds. For Go trials, each trait was paired with a delayed Go cue that required participants to press a button as quickly as possible before the trait’s offset. To maintain participants’ attention, we used an adaptive response window. Specifically, the go-signal-delay (GSD, the delay between trait onset and Go-cue onset) was approximately 0.9 second. If the participants gave a timely response (i.e., button press before the offset of the trait), the GSD was increased by 17 ms to increase task difficulty. If participants failed to make a button press before the offset of the trial, the GSD was reduced by 50 ms to reduce task difficulty (Salomon et al., 2018; Schonberg et al., 2014). Conversely, for NoGo trials, participants merely viewed and listened to the traits without any behavioral responses. All 60 positive traits were presented randomly in each of the five blocks during the CAT, resulting in a total of 300 trials. Participants could take a 0.5-1-minute break between blocks. While previous CAT research adopts over 10 blocks of training (Salomon et al., 2018), we chose to only include 5 blocks so as to avoid ceiling effect in subsequent memory recall. This CAT task was followed by a 5-minute working memory task, serving as distractions.

Following the working memory task, participants proceeded to a 3-minute post-CAT self-referential free recall task, which was identical to the baseline task. Subsequently, a post-CAT probe task was administered to evaluate the impact of CAT.

In the probe task (see Figure 1D), participants were presented with Go and NoGo traits in pairs and were asked to choose which trait would be more self-descriptive. Within each trial, the Go and NoGo traits were matched on baseline endorsement ratings, so that preferential choices of Go traits would indicate the CAT training effects. The positions of the Go/NoGo traits per pair were randomly assigned to the upper-left/right or upper-right/left sides of the monitor in the first block, and were swapped in the second block. Each trial started with a fixation cross (1 second), followed by the side-by-side presentation of two traits. Participants selected the trait that would best describe them by clicking a push button below the trait within 2.5 seconds. The chosen trait was then highlighted by a button-press shaped image for 0.5 seconds. If participants exceeded the 4-second response time, a prompt would appear during the confirmation phase, urging them to respond quickly. We excluded trials with response times exceeding 3 seconds, accounting for potential mouse delays.

#### Nap targeted memory reactivation (TMR)

Participants took a 90-minute nap in a quiet, darkened sleep chamber. Background white noise (at ∼38 dB) was played to participants throughout the duration of the nap via a loudspeaker placed near the bed. Participants’ brain and physiological activities were continuously monitored during the map. Upon participants entered SWS for at least 2 minutes, we presented spoken positive traits (the same spoken traits presented during the SRET and CAT tasks) at approximately 40 dB. Note that the spoken traits (∼40 dB) were played against the background white noise (∼38 dB), yet remained subtle to avoid arousal and waking participants up.

The TMR began with playing a neutral trait (∼0.6 s) for three times, ensuring that the auditory stimulation would not wake participants up. We started playing the spoken traits if participants did not show signs of arousal or changes in NREM sleep stage. During each round of the TMR, half of the positive Go traits (i.e., 15 traits) were played together with the neutral trait as a control word. Each trait last for about 1 second, with a randomized interstimulus interval of 5–6 seconds. TMR continued as participants remained in the SWS, with a minimal repetition of three rounds of stimulation, resulting in at least 3 X 16 = 48 trials for TMR-related EEG analyses.

Specifically, participants were exposed to spoken traits once they entered a sustained SWS period lasting at least 2 minutes. The TMR procedure was discontinued after 30 minutes, or earlier if EEG recordings indicated micro-arousal or full awakening. If no SWS was detected within the first 40 minutes, the presentation of spoken traits commenced during the N2 sleep stage. After a total sleep session of 90 minutes, participants were awakened if they were in the N1 or N2 sleep stages, or we waited until they transitioned to these stages before awakening them. A brief 5-minute break was provided upon awakening to mitigate the effects of sleep inertia.

#### Post-TMR tests

Participants completed the self-referential free call task, probe task, and SRET task. Here, the probe task instructions were identical to the post-CAT probe task, but with randomized ‘Go’ and ‘NoGo’ trait positions. The SRET was similar to the baseline SRET except that participants only made a Yes/No binary response to each trait, omitting the rating part.

#### One-week delayed tests

Participants returned to the lab about one week later to complete the delayed tests in the following order: (1) a 3-minute self-referential free call task; (2) a probe task; (3) a SRET task. The tasks were identical to the tasks in the post-TMR. Participants were not informed of the delayed tasks ahead of the time. Upon completing all tasks, participants were debriefed and paid.

## Quantification and statistical analysis

### Behavioral data analysis

Statistical analyses were carried out using R (Version 4.2.1., R Core Team (2020). We performed (G)LMMs fitted via ‘*glmer’* and ‘*lmer’* functions of the ‘*lme4’* R package (Bates et al. 2014 June 23) to analyze the CAT- and TMR-induced behavioral changes. For statistical significance testing, we used Type III Analysis of Variance with the Satterthwaite approximation method for the LMM and Type II Wald Chi-Square tests for the GLMM. We followed up significant effects with post-hoc comparisons in *emmeans* (Lenth et al., 2022) to derive the estimated marginal means from each model. Model predictions were visualized with the ‘*plot_model*’ function from the *sjPlot* package (Lüdecke, 2023). Unless otherwise stated, we used the False Discovery Rate (FDR) method to adjust for multiple comparisons to control for false-positive results. The significance threshold (alpha level) was set at 0.05.

#### Self-referential preference choices in the probe task

Following previous CAT research (Botvinik-Nezer et al., 2020; Salomon et al., 2018; Schonberg et al., 2014), we ran generalized linear mixed models (GLMMs) to compare the odds of choosing Go traits against the chance level (50%, log odds = 0; odds ratio = 1) during post-CAT phase. Given the alternation of Go/NoGo positions (left and right) in two blocks, we included Go position as a covariate in our model. The GLMM was defined as:

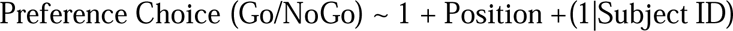

#### Self-referential endorsement in the SRET

We employed a generalized linear mixed model (GLMM) to examine how TMR conditions (Go-cued, Go-uncued, and NoGo-uncued) influenced participants’ endorsement for positive traits across time (baseline, post-TMR and delay). We used baseline endorsement rating as a covariate and participant as random effect. The model was defined as:

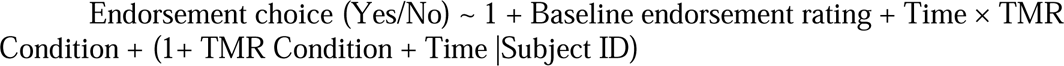

Subsequently, we employed a linear mixed model (LMM), incorporating the same factors as used in the preceding GLMM for binary choice outcomes to analyze RTs when endorsing positive traits except that we removed TMR as random slope due to singular fitting:

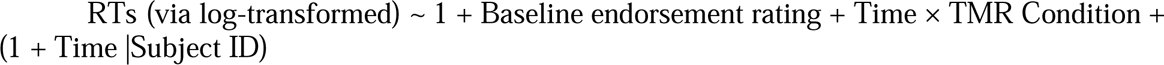

In addition, we also ran a GLMM to examine the endorsement changes for negative traits, using time as a fixed effect:

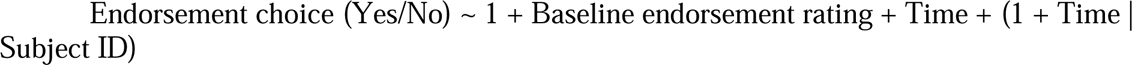

Lastly, in order to assess whether CAT alone influenced the response speed during the endorsement of positive traits, we employed another LMM in two additional behavioral samples. These samples comprised one group that underwent only CAT training (referred to as the ‘active’ group) and another group that received no CAT training (referred to as the ‘passive’ group). Detailed information on these two behavioral samples can be found in the Supplementary Online Material (SOM). The LMM incorporated several fixed effects: group (active vs. passive), time (baseline, post-CAT, delay), and CAT (Go vs. NoGo). Additionally, the baseline endorsement rating was included as a covariate. The model also accounted for random effects at the participant level:

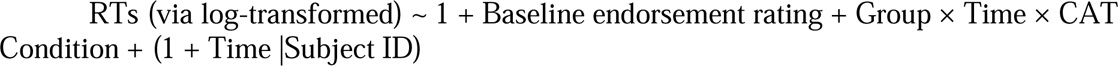

#### Self-referential memories in the free recall task

To understand how TMR affect self-referential memories across time, we ran a GLMM using TMR (Go-cued, Go-uncued, and NoGo-uncued), and time (baseline, post-CAT, post-TMR, and delay) as fixed effects, baseline endorsement rating as covariate, participant as random effect. Time was removed from random slope given singular fitting issue. The model was defined as follows:

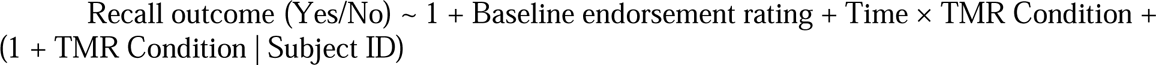

Additionally, a GLMM was applied to analyze negative traits, using time as a fixed effect:

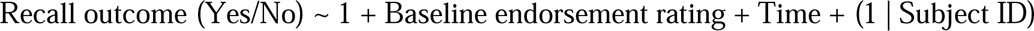

Finally, incorporating data from two additional samples — one with only CAT training and another with no training (see SOM for details regarding behavioral samples)— we expanded our analysis to encompass three distinct groups. To assess delayed recall across these groups, we employed a GLMM on delayed recall performance with baseline and post-CAT recall as covariate, training groups (i.e., both TMR and CAT trained, only CAT trained, no CAT trained) as fixed effect:

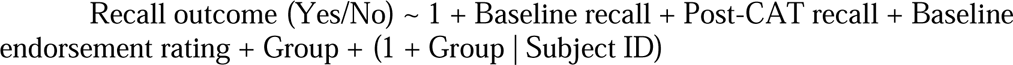

### EEG data analysis

#### EEG data pre-processing

Continuous EEGs were recorded using a 63-channel customized cap with passive Ag/AgCl electrodes via a BrainAmp amplifier with a 1000 Hz sampling rate (Brain Products, Gilching, Germany). Electrodes were positioned according to the International 10–10 system. The ground electrode was located at AFz, with FCz as the on-line reference electrode. Impedances were kept below 20 kΩ. We placed one electro-oculography (EOG) electrode under participants’ left eyes and bipolar electromyography (EMG) electrodes on their chins to monitor eye movements and muscle activity during sleep.

EEG data were pre-processed using custom-written scripts and the MATLAB Toolbox EEGLAB (Delorme & Makeig, 2004). First, nap EEG data were down-sampled to 250 Hz, notch-filtered at 50 Hz, and then re-referenced to the averaged mastoids. Second, EEG data were band-pass filtered at 0.5 to 40 Hz. While EOG and EMG data were used for sleep staging, these data were not used in the time-frequency analysis.

#### Offline Sleep Stage Scoring

Sleep stages, including N1, N2, Slow-Wave Sleep (SWS), and Rapid Eye Movement (REM), were scored using EEG (Channel C4), EOG, and EMG patterns. This process employed algorithms from the YASA open-source Python Toolbox (Vallat & Walker, 2021). Consistent with YASA guidelines, the EEG data were initially re-referenced to FPz before conducting the staging analysis. Table 1 presents the sleep staging results for 34 participants (One participants only reserved 28 min EEG data including TMR stage).

**Table 1.** Sleep stages parameters (meanLJ±LJSEM, in minutes, N = 34).

| Total time | Wake | N1 | N2 | N3 | REM |
| --- | --- | --- | --- | --- | --- |
| 90.10 | 7.76 | 7.18 | 36.00 | 27.00 | 12.36 |
| ± | ± | ± | ± | ± | ± |
| 1.27 | 1.19 | 0.63 | 1.56 | 1.83 | 1.27 |

#### EEG time-frequency analysis

Before analyzing cue-elicited time-frequency EEG power changes, the cue-elicited EEG data were epoched into −1.5 to 5.5 second segments, relative to the onset of each cued trait word. This long epoch ensured that we had enough edge artifact-free segments for each clean epoch to assess TMR benefits (−1 to 3 seconds). Epochs with artefacts were visually inspected and removed. Time-frequency decomposition was performed in the Fieldtrip open-source MATLAB toolbox (Oostenveld et al., 2011). We used 3 to 10 cycles in a step of 0.5 Hz Morlet wavelet and baseline corrected using z-transformation of all trials from −1 to −0.1 seconds relative to the cue onset. Following previous sleep and TMR studies (Mölle et al., 2011; Wilhelm et al., 2020; Xia et al., 2023; Züst et al., 2019), we calculated the mean EEG power over frontal-central channel (F1/2, FC1/2, C1/2, Fz, Cz) to ensure the robustness of results. The calculated time-frequency decompositions were then down-sampled to 50 Hz. To investigate cue-elicited EEG activity, we employed the rigorous cluster-based one-sample permutation t-test (cluster-thresholding *p* at 0.001) to identify the significant cluster against zero across all participants in the time-frequency domain (Maris & Oostenveld, 2007).

#### Traveling wave analysis

We employed a traveling wave analysis approach similar to that used by Alamia et al. 2023. This involved calculating values for spontaneous slow backward (from the anterior to posterior regions) and forward (from posterior to anterior regions) traveling waves using 2D Fast Fourier Transform (FFT) on the time-electrode EEG signals. The power measured in the upper right and lower right quadrants corresponded to the amount of backward and forward propagating waves, respectively, as shown in Figure 4A. Specifically, we used EEG data from the interval after cue onset [0, 2000] ms during NREM stage 3 (based on YAS staging result) across midline positions (POz, Pz, CPz, Cz, Fz, FPz) to create time-electrode EEG representations. To establish a baseline, we shuffled the electrodes and repeated this process. For the slow wave range (1–4 Hz), we identified the maximum values in the 2D-FFT spectra in both the actual data (BW and FW) and the shuffled data (*BWss* and *FWss*). The magnitude of the backward and forward traveling waves, expressed in decibels [dB], was calculated as:

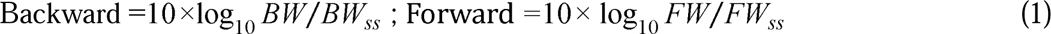

#### Brain-behavior association analysis

To establish a direct link between TMR-induced behavioral changes and TMR-elicited EEG activity, we extracted the averaged power within the identified significant positive clusters, and also calculated mean traveling waves for each participant at each item level. Then we performed a series of LMMs using EEG power and traveling waves to predict post-TMR SRET performance metrics, including endorsement choices and positive endorsement RTs. All EEG metrics were centered before being included as fixed effects. We used GLMM to predict endorsement choice (Yes/No) and LMMs to predict RTs for endorsing positive traits. The models were defined as:

1. Post-TMR endorsement choice (Yes/No) ∼ 1 + Positive delta-theta-alpha cluster/ Positive sigma-beta cluster/ Forward traveling wave / Backward traveling wave + Baseline Choice + Repetition + (1|Subject ID).
2. Delay endorsement choice (Yes/No) ∼ 1 + Positive delta-theta-alpha cluster/ Positive sigma-beta cluster/ Forward traveling wave / Backward traveling wave + Baseline Choice + Repetition + (1|Subject ID).
3. Post-TMR RTs for endorsing positive traits ∼ 1 Positive delta-theta-alpha cluster/ Positive sigma-beta cluster/ Forward traveling wave / Backward traveling wave + Baseline RTs + Repetition + (1|Subject ID).
4. Delay RTs for endorsing positive traits ∼ 1 + Positive delta-theta-alpha cluster/ Positive sigma-beta cluster/ Forward traveling wave / Backward traveling wave + Baseline RTs + Repetition + (1|Subject ID).

## References

Abdellahi, M. E. A., Koopman, A. C. M., Treder, M. S., & Lewis, P. A. (2023). Targeting targeted memory reactivation: Characteristics of cued reactivation in sleep. NeuroImage, 266, 119820. 10.1016/j.neuroimage.2022.119820

Adamantidis, A. R., Gutierrez Herrera, C., & Gent, T. C. (2019). Oscillating circuitries in the sleeping brain. Nature Reviews Neuroscience, 20(12), Article 12. 10.1038/s41583-019-0223-4

Ai, S., Yin, Y., Chen, Y., Wang, C., Sun, Y., Tang, X., Lu, L., Zhu, L., & Shi, J. (2018). Promoting subjective preferences in simple economic choices during nap. eLife, 7, e40583. 10.7554/eLife.40583

Alamia, A., Terral, L., D’ambra, M. R., & VanRullen, R. (2023). Distinct roles of forward and backward alpha-band waves in spatial visual attention. eLife, 12, e85035. 10.7554/eLife.85035

Antony, J. W., Cheng, L. Y., Brooks, P. P., Paller, K. A., & Norman, K. A. (2018). Competitive learning modulates memory consolidation during sleep. Neurobiology of Learning and Memory, 155, 216–230. 10.1016/j.nlm.2018.08.007

Antony, J. W., Ferreira, C. S., Norman, K. A., & Wimber, M. (2017). Retrieval as a Fast Route to Memory Consolidation. Trends in Cognitive Sciences, 21(8), 573–576. 10.1016/j.tics.2017.05.001

Barner, C., Werner, A.-S., Schörk, S., Born, J., & Diekelmann, S. (2023). The effects of sleep and targeted memory reactivation on the consolidation of relevant and irrelevant information. Frontiers in Sleep, 2. https://www.frontiersin.org/articles/10.3389/frsle.2023.1187170

Bates, D., Mächler, M., Bolker, B., & Walker, S. (2014). Fitting Linear Mixed-Effects Models using lme4. 10.48550/arXiv.1406.5823

Batterink, L. J., Westerberg, C. E., & Paller, K. A. (2017). Vocabulary learning benefits from REM after slow-wave sleep. Neurobiology of Learning and Memory, 144, 102–113. 10.1016/j.nlm.2017.07.001

Beck, A. T., Steer, R. A., Ball, R., & Ranieri, W. F. (1996). Comparison of Beck Depression Inventories-IA and-II in Psychiatric Outpatients. Journal of Personality Assessment, 67(3), 588–597. 10.1207/s15327752jpa6703_13

Blume, C., del Giudice, R., Lechinger, J., Wislowska, M., Heib, D. P. J., Hoedlmoser, K., & Schabus, M. (2017). Preferential processing of emotionally and self-relevant stimuli persists in unconscious N2 sleep. Brain and Language, 167, 72–82. 10.1016/j.bandl.2016.02.004

Born, J., & Wilhelm, I. (2012). System consolidation of memory during sleep. Psychological Research, 76(2), 192–203. 10.1007/s00426-011-0335-6

Botvinik-Nezer, R., Salomon, T., & Schonberg, T. (2020). Enhanced Bottom-Up and Reduced Top-Down fMRI Activity Is Related to Long-Lasting Nonreinforced Behavioral Change. Cerebral Cortex, 30(3), 858–874. 10.1093/cercor/bhz132

Brodt, S., Inostroza, M., Niethard, N., & Born, J. (2023). Sleep—A brain-state serving systems memory consolidation. Neuron, 111(7), 1050–1075. 10.1016/j.neuron.2023.03.005

Cairney, S. A., Lindsay, S., Sobczak, J. M., Paller, K. A., & Gaskell, M. G. (2016). The Benefits of Targeted Memory Reactivation for Consolidation in Sleep are Contingent on Memory Accuracy and Direct Cue-Memory Associations. Sleep, 39(5), 1139–1150. 10.5665/sleep.5772

Collins, A. C., & Winer, E. S. (2023). Self-Referential Processing and Depression: A Systematic Review and Meta-Analysis. Clinical Psychological Science, 21677026231190390. 10.1177/21677026231190390

Colombo, D., Fernández-Álvarez, J., Suso-Ribera, C., Cipresso, P., García-Palacios, A., Riva, G., & Botella, C. (2020). Biased Affective Forecasting: A Potential Mechanism That Enhances Resilience and Well-Being. Frontiers in Psychology, 11. https://www.frontiersin.org/article/10.3389/fpsyg.2020.01333

Creery, J. D., Oudiette, D., Antony, J. W., & Paller, K. A. (2015). Targeted Memory Reactivation during Sleep Depends on Prior Learning. Sleep, 38(5), 755–763. 10.5665/sleep.4670

Dainer-Best, J., Lee, H. Y., Shumake, J. D., Yeager, D. S., & Beevers, C. G. (2018). Determining optimal parameters of the self-referent encoding task: A large-scale examination of self-referent cognition and depression. Psychological Assessment, 30(11), 1527–1540. 10.1037/pas0000602

Dainer-Best, J., Trujillo, L. T., Schnyer, D. M., & Beevers, C. G. (2017). Sustained engagement of attention is associated with increased negative self-referent processing in major depressive disorder. Biological Psychology, 129, 231–241. 10.1016/j.biopsycho.2017.09.005

Delorme, A., & Makeig, S. (2004). EEGLAB: An open source toolbox for analysis of single-trial EEG dynamics including independent component analysis. Journal of Neuroscience Methods, 134(1), 9–21. 10.1016/j.jneumeth.2003.10.009

Denis, D., & Payne, J. D. (2023). Targeted memory reactivation during non-rapid eye movement sleep enhances neutral, but not negative, components of memory. bioRxiv. 10.1101/2023.05.26.542120

Derry, P. A., & Kuiper, N. A. (1981). Schematic processing and self-reference in clinical depression. Journal of Abnormal Psychology, 90(4), 286–297. 10.1037/0021-843X.90.4.286

Diekelmann, S., & Born, J. (2010). The memory function of sleep. Nature Reviews Neuroscience, 11(2). 10.1038/nrn2762

Göldi, M., van Poppel, E. A. M., Rasch, B., & Schreiner, T. (2019). Increased neuronal signatures of targeted memory reactivation during slow-wave up states. Scientific Reports, 9(1). 10.1038/s41598-019-39178-2

Guenther, C. L., & Alicke, M. D. (2010). Deconstructing the better-than-average effect. Journal of Personality and Social Psychology, 99(5), 755–770. https://psycnet.apa.org/doi/10.1037/a0020959

Guttesen, A. á V., Gaskell, M. G., & Cairney, S. A. (2023). Delineating memory reactivation in sleep with verbal and non-verbal retrieval cues. bioRxiv. 10.1101/2023.03.02.530762

Halgren, M., Ulbert, I., Bastuji, H., Fabó, D., Erőss, L., Rey, M., Devinsky, O., Doyle, W. K., Mak-McCully, R., Halgren, E., Wittner, L., Chauvel, P., Heit, G., Eskandar, E., Mandell, A., & Cash, S. S. (2019). The generation and propagation of the human alpha rhythm. Proceedings of the National Academy of Sciences, 116(47), 23772–23782. 10.1073/pnas.1913092116

Hangya, B., Tihanyi, B. T., Entz, L., Fabó, D., Eró ss, L., Wittner, L., Jakus, R., Varga, V., Freund, T. F., & Ulbert, I. (2011). Complex Propagation Patterns Characterize Human Cortical Activity during Slow-Wave Sleep. Journal of Neuroscience, 31(24). 10.1523/JNEUROSCI.1498-11.2011

Hobbs, C., Sui, J., Kessler, D., Munafò, M. R., & Button, K. S. (2023). Self-processing in relation to emotion and reward processing in depression. Psychological Medicine, 53(5), 1924–1936. 10.1017/S0033291721003597

Hoffmann, J., Hobbs, C., Moutoussis, M., & Button, K. (2023). Lack of optimistic biases in depression and social anxiety is reflected in reduced positive self-beliefs, but distinct processing of social feedback. PsyArXiv. 10.31234/osf.io/h6ety

Hu, X., Antony, J. W., Creery, J. D., Vargas, I. M., Bodenhausen, G. V., & Paller, K. A. (2015). Unlearning implicit social biases during sleep. Science, 348(6238), 1013–1015. 10.1126/science.aaa3841

Hu, X., Cheng, L. Y., Chiu, M. H., & Paller, K. A. (2020). Promoting memory consolidation during sleep: A meta-analysis of targeted memory reactivation. Psychological Bulletin, 146(3), 218–244. 10.1037/bul0000223

Hutchison, I. C., Pezzoli, S., Tsimpanouli, M.-E., Abdellahi, M. E. A., Pobric, G., Hulleman, J., & Lewis, P. A. (2021). Targeted memory reactivation in REM but not SWS selectively reduces arousal responses. Communications Biology, 4(1). 10.1038/s42003-021-01854-3

Itzkovitch, A., Bar Or, M., & Schonberg, T. (2022). Cue-approach training for food behavior. Current Opinion in Behavioral Sciences, 47, 101202. 10.1016/j.cobeha.2022.101202

John, O. P., Donahue, E. M., & Kentle, R. L. (1991). The Big Five Inventory—Versions 4a and 54 (Vol. 10). Berkeley, CA: University of California, Berkeley, Institute of Personality and Social Research.

Klinzing, J. G., Niethard, N., & Born, J. (2019). Mechanisms of systems memory consolidation during sleep. Nature Neuroscience, 22(10). 10.1038/s41593-019-0467-3

Konovalov, A., & Krajbich, I. (2019). Revealed strength of preference: Inference from response times. Judgment and Decision Making, 14(4), 381–394. 10.1017/S1930297500006082

Kurth, S., Riedner, B. A., Dean, D. C., O’Muircheartaigh, J., Huber, R., Jenni, O. G., Deoni, S. C. L., & LeBourgeois, M. K. (2017). Traveling Slow Oscillations During Sleep: A Marker of Brain Connectivity in Childhood. Sleep, 40(9). 10.1093/sleep/zsx121

Lehmann, M., Schreiner, T., Seifritz, E., & Rasch, B. (2016). Emotional arousal modulates oscillatory correlates of targeted memory reactivation during NREM, but not REM sleep. Scientific Reports, 6(1). 10.1038/srep39229

Lenth, R. V., Buerkner, P., Herve, M., Love, J., Miguez, F., Riebl, H., & Singmann, H. (2022). emmeans: Estimated Marginal Means, aka Least-Squares Means (1.7.2) [Computer software]. https://CRAN.R-project.org/package=emmeans

Lewis, P. A., & Bendor, D. (2019). How Targeted Memory Reactivation Promotes the Selective Strengthening of Memories in Sleep. Current Biology, 29(18), R906–R912. 10.1016/j.cub.2019.08.019

Liu, J., Xia, T., Chen, D., Yao, Z., Zhu, M., Antony, J. W., Lee, T. M. C., & Hu, X. (2023). Item-specific neural representations during human sleep support long-term memory. PLOS Biology, 21(11), e3002399. 10.1371/journal.pbio.3002399

Lou, Y., Lei, Y., Mei, Y., Leppänen, P. H. T., & Li, H. (2019). Review of Abnormal Self-Knowledge in Major Depressive Disorder. Frontiers in Psychiatry, 10. 10.3389/fpsyt.2019.00130

Lüdecke, D. (2021). sjPlot: data visualization for statistics in social science (2.8. 13) [Computer software]. https://CRAN.R-project.org/package=sjPlot

MacDonald, K. J., & Cote, K. A. (2021). Contributions of post-learning REM and NREM sleep to memory retrieval. Sleep Medicine Reviews, 59, 101453. 10.1016/j.smrv.2021.101453

Maris, E., & Oostenveld, R. (2007). Nonparametric statistical testing of EEG- and MEG-data. Journal of Neuroscience Methods, 164(1), 177–190. 10.1016/j.jneumeth.2007.03.024

Massimini, M., Huber, R., Ferrarelli, F., Hill, S., & Tononi, G. (2004). The Sleep Slow Oscillation as a Traveling Wave. Journal of Neuroscience, 24(31), Article 31. 10.1523/JNEUROSCI.1318-04.2004

Mölle, M., Bergmann, T. O., Marshall, L., & Born, J. (2011). Fast and Slow Spindles during the Sleep Slow Oscillation: Disparate Coalescence and Engagement in Memory Processing. Sleep, 34(10), 1411–1421. 10.5665/SLEEP.1290

Muller, L., Chavane, F., Reynolds, J., & Sejnowski, T. J. (2018). Cortical travelling waves: Mechanisms and computational principles. Nature Reviews Neuroscience, 19(5), 255–268. 10.1038/nrn.2018.20

Murphy, M., Riedner, B. A., Huber, R., Massimini, M., Ferrarelli, F., & Tononi, G. (2009). Source modeling sleep slow waves. Proceedings of the National Academy of Sciences, 106(5), Article 5. 10.1073/pnas.0807933106

Oostenveld, R., Fries, P., Maris, E., & Schoffelen, J.-M. (2011). FieldTrip: Open Source Software for Advanced Analysis of MEG, EEG, and Invasive Electrophysiological Data. Computational Intelligence and Neuroscience, 2011, 1–9. 10.1155/2011/156869

Orth, U., Robins, R. W., & Link to external site, this link will open in a new window. (2022). Is high self-esteem beneficial? Revisiting a classic question. American Psychologist, 77(1), 5–17. 10.1037/amp0000922

Oudiette, D., Antony, J. W., Creery, J. D., & Paller, K. A. (2013). The role of memory reactivation during wakefulness and sleep in determining which memories endure. Journal of Neuroscience, 33(15), 6672–6678. 10.1523/JNEUROSCI.5497-12.2013

Oudiette, D., & Paller, K. A. (2013). Upgrading the sleeping brain with targeted memory reactivation. Trends in Cognitive Sciences, 17(3), 142–149. 10.1016/j.tics.2013.01.006

Patton, J. H., Stanford, M. S., & Barratt, E. S. (1995). Factor structure of the Barratt Impulsiveness Scale. Journal of Clinical Psychology, 51(6), 768–774. 10.1002/1097-4679(199511)51:6<768::AID-JCLP2270510607>3.0.CO;2-1

Petzka, M., Chatburn, A., Charest, I., Balanos, G. M., & Staresina, B. P. (2022). Sleep spindles track cortical learning patterns for memory consolidation. Current Biology, 32(11), 2349–2356.e4. 10.1016/j.cub.2022.04.045

R Core Team (2020). (n.d.). R: a language and environment for statistical computing. R Foundation for Statistical Computing (4.1.3) [Computer software]. https://www.r-project.org/index.html

Rakowska, M., Abdellahi, M. E. A., Bagrowska, P., Navarrete, M., & Lewis, P. A. (2021). Long term effects of cueing procedural memory reactivation during NREM sleep. NeuroImage, 244, 118573. 10.1016/j.neuroimage.2021.118573

Rasch, B., & Born, J. (2013). About Sleep’s Role in Memory. Physiological Reviews, 93(2), 681–766. 10.1152/physrev.00032.2012

Raskin, R., & Hall, C. S. (1981). The Narcissistic Personality Inventory: Alternative Form Reliability and Further Evidence of Construct Validity. Journal of Personality Assessment, 45(2), 159. 10.1207/s15327752jpa4502_10

Romero, N., Sanchez, A., Vázquez, C., & Valiente, C. (2016). Explicit self-esteem mediates the relationship between implicit self-esteem and memory biases in major depression. Psychiatry Research, 242, 336–344. 10.1016/j.psychres.2016.06.003

Rosenberg, M. (1965). Society and the adolescent self-image. Princeton, NJ: Princeton University Press.

Salomon, T., Botvinik-Nezer, R., Gutentag, T., Gera, R., Iwanir, R., Tamir, M., & Schonberg, T. (2018). The Cue-Approach Task as a General Mechanism for Long-Term Non-Reinforced Behavioral Change. Scientific Reports, 8(1), 3614. 10.1038/s41598-018-21774-3

Schechtman, E., Antony, J. W., Lampe, A., Wilson, B. J., Norman, K. A., & Paller, K. A. (2021). Multiple memories can be simultaneously reactivated during sleep as effectively as a single memory. Communications Biology, 4(1), 1–13. 10.1038/s42003-020-01512-0

Schechtman, E., Heilberg, J., & Paller, K. A. (2023). Memory consolidation during sleep involves context reinstatement in humans. Cell Reports, 42(4), 112331. 10.1016/j.celrep.2023.112331

Schonberg, T., Bakkour, A., Hover, A. M., Mumford, J. A., Nagar, L., Perez, J., & Poldrack, R. A. (2014). Changing value through cued approach: An automatic mechanism of behavior change. Nature Neuroscience, 17(4), 625–630. 10.1038/nn.3673

Schonberg, T., & Katz, L. N. (2020). A Neural Pathway for Nonreinforced Preference Change. Trends in Cognitive Sciences, 24(7), 504–514. 10.1016/j.tics.2020.04.002

Schreiner, T., Kaufmann, E., Noachtar, S., Mehrkens, J.-H., & Staudigl, T. (2022). The human thalamus orchestrates neocortical oscillations during NREM sleep. Nature Communications, 13(1), Article 1. 10.1038/s41467-022-32840-w

Schreiner, T., Lehmann, M., & Rasch, B. (2015). Auditory feedback blocks memory benefits of cueing during sleep. Nature Communications, 6(1), Article 1. 10.1038/ncomms9729

Schreiner, T., Petzka, M., Staudigl, T., & Staresina, B. P. (2021). Endogenous memory reactivation during sleep in humans is clocked by slow oscillation-spindle complexes. Nature Communications, 12(1), Article 1. 10.1038/s41467-021-23520-2

Schreiner, T., & Rasch, B. (2015). Boosting Vocabulary Learning by Verbal Cueing During Sleep. Cerebral Cortex, 25(11), 4169–4179. 10.1093/cercor/bhu139

Sowislo, J. F. & Orth, U. (2013). Does low self-esteem predict depression and anxiety? A meta-analysis of longitudinal studies. Psychological Bulletin, 139(1), 213–240. 10.1037/a0028931

Spielberger, C. D. (1983). State-Trait Anxiety Inventory, Form Y (STAI). Palo Alto, CA: Consulting Psychologicals Press. https://journal.sipsych.org/index.php/IJP/article/view/620

Taylor, S. E., & Brown, J. D. (1988). Illusion and well-being: A social psychological perspective on mental health. Psychological Bulletin, 103(2), 193–210. 10.1037/0033-2909.103.2.193

Vallat, R., & Walker, M. P. (2021). An open-source, high-performance tool for automated sleep staging. eLife, 10, e70092. 10.7554/eLife.70092

Walker, M. P., & Stickgold, R. (2006). Sleep, Memory, and Plasticity. Annual Review of Psychology, 57(1), 139–166. 10.1146/annurev.psych.56.091103.070307

Watson, L. A., Dritschel, B., Obonsawin, M. C., & Jentzsch, I. (2007). Seeing yourself in a positive light: Brain correlates of the self-positivity bias. Brain Research, 1152, 106–110. 10.1016/j.brainres.2007.03.049

Weisenburger, R. L., Dainer-Best, J., Zisser, M., McNamara, M., & Beevers, C. G. (2023). Negative Self-Referent Cognition Predicts Future Depression Symptom Change: An Intensive Sampling Approach. PsyArXiv. 10.31234/osf.io/3tn4c

Wilhelm, I., Schreiner, T., Beck, J., & Rasch, B. (2020). No effect of targeted memory reactivation during sleep on retention of vocabulary in adolescents. Scientific Reports, 10(1), 1–9. 10.1038/s41598-020-61183-z

Wisco, B. E. (2009). Depressive cognition: Self-reference and depth of processing. Clinical Psychology Review, 29(4), 382–392. 10.1016/j.cpr.2009.03.003

Xia, T., Yao, Z., Guo, X., Liu, J., Chen, D., Liu, Q., Paller, K. A., & Hu, X. (2023). Updating memories of unwanted emotions during human sleep. Current Biology, 33(2), 309–320.e5. 10.1016/j.cub.2022.12.004

Zell, E., Strickhouser, J. E., Sedikides, C., & Alicke, M. D. (2020). The better-than-average effect in comparative self-evaluation: A comprehensive review and meta-analysis. Psychological Bulletin, 146(2), 118–149. 10.1037/bul0000218

Zhang, H., Watrous, A. J., Patel, A., & Jacobs, J. (2018). Theta and Alpha Oscillations Are Traveling Waves in the Human Neocortex. Neuron, 98(6), 1269–1281.e4. 10.1016/j.neuron.2018.05.019

Züst, M. A., Ruch, S., Wiest, R., & Henke, K. (2019). Implicit Vocabulary Learning during Sleep Is Bound to Slow-Wave Peaks. Current Biology, 29(4), 541–553.e7. 10.1016/j.cub.2018.12.038

