## Supplementary material for "Reactivating Positive Personality Traits During Sleep Promotes Positive Self-Referential Processing": SOM

### Baseline endorsement rating for Go versus NoGo traits

We ran a one-sample paired t-test analysis on ratings for Go and NoGo traits to confirm that there was no significant difference of rating. Result revealed no significant difference between Go and NoGo trait ratings (*t* (34) = 0.55, *p* = 0.586).

### Participants from two behavioural samples

In this behavioral sample, we recruited 74 participants from Shenzhen University who were compensated monetarily at a rate of 50 RMB per hour (appropriately 7.8 USD). Participants were randomly assigned to either the active- or passive-CAT group. The active CAT was similar like in the CAT in the main text. The only difference between the active- and passive-CAT group is that participants in the passive-CAT group were only instructed to look at the words presented on the screen and were not required to press a button when the white cue was displayed.

We excluded four participants (two from each group) due to lack of attention during CAT, leaving 34 participants in each group for analysis (active-CAT group: 17 Males, Mage ± SD = 22.57 ± 2.25; passive-CAT group, 18 Males, Mage ± SD = 22.71 ± 1.60). We only included participants with normal or corrected-to-normal vision, no history of mental illness or neurological disorder, and no current or history of sleeping disorders. The local ethics committee approved the study, and all participants signed the consent form before participating the experiment. Only data from self-referential encoding task were analyzed in this paper.

**Figure S1**

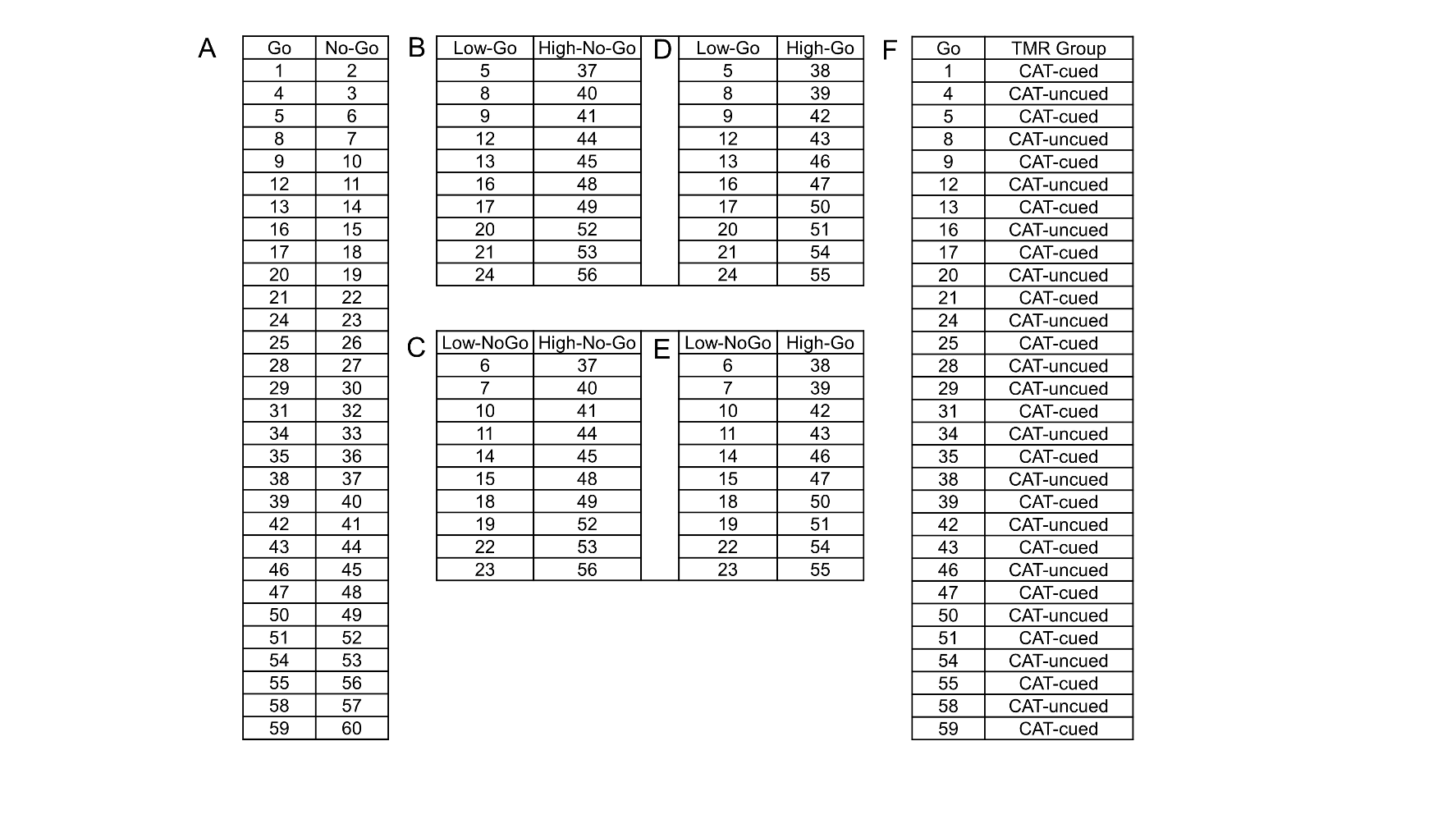

*Sorting and Pair-Matching Procedure in TMR experiment.*

1A. All trait words were ranked from lowest (1) to highest (60) based on their initial endorsement rating in the baseline SRET. Go and NoGo items of the same rank were paired, generating 30 pairs in total, ensuring similar initial endorsement ratings between Go and NoGo items in each pair. Trait words were marked as low-rating (ranks 5:24) and high-rating (ranks 37:56), with 10 unique pairs in each low- and high-rating categories. 1B~E. Control pairings were created to rule out mere exposure effects and avoid suspicions of pairing rules, including low-value Go with high-value NoGo (S1B), low-value NoGo with high-value NoGo (S1C), low-value Go with high-value Go (S1C), and low-value NoGo with high-value Go (S1E), but these pairings were not analysed. 1F. Go traits were grouped into to Go-cued and Go-uncued categories using a counterbalanced order.

### Table S1 *Descriptive statistics of questionnaire scores in TMR experiment.*

| Questionnaire Mean S.D. | | |
| --- | --- | --- |
| Rosenberg Self-Esteem Scale (RSES) | 15.743 | 1.686 |
| Depression symptoms, BDI-II | 7.686 | 6.356 |
| Barratt Impulsiveness Scale (BIS) | 57.286 | 8.234 |
| Narcissistic Personality Inventory (NPI) | 24.286 | 1.545 |
| Trait anxiety, STAI | 40.314 | 7.336 |
| State anxiety, STAI | 36.743 | 7.406 |
| Extraversion, BFI | 24.371 | 5.331 |
| Agreeableness, BFI | 33.886 | 4.813 |
| Conscientiousness, BFI | 27.857 | 4.864 |
| Neuroticism, BFI | 22.6 | 5.077 |
| Openness, BFI | 36.171 | 5.377 |

*Note.* CAT = Cue approach training. BFI = Big Five Inventory

### A Complete List of Personality Adjectives

The translated version of the document is available as supplementary material on our OSF repository: <https://osf.io/rztdh/?view_only=1f6596bd31a049199a5df4d4d7c764c4>

| English Version | Valence Rating | Arousal Rating | Familiarity Rating |
| --- | --- | --- | --- |
| optimistic | 7.55 | 4.8 | 7.35 |
| companionable | 7.5 | 4.85 | 7.3 |
| considerate | 7.6 | 4.5 | 7.15 |
| courageous | 7.3 | 6.8 | 6.9 |
| diligent | 6.7 | 4.9 | 7 |
| innocent | 6.25 | 3.85 | 7.2 |
| educated | 7 | 3.55 | 6.25 |
| friendly | 7.55 | 4.6 | 7.5 |
| reliable | 7.5 | 3.95 | 6.95 |
| good-natured | 7.4 | 4.3 | 6.65 |
| gracious | 7.35 | 3.75 | 6.8 |
| kind | 7.4 | 3.9 | 7.6 |
| persistent | 6.6 | 4.95 | 7.05 |
| candid | 6.6 | 4.1 | 6.65 |
| outgoing | 6.35 | 5.55 | 6.65 |
| curious | 6.85 | 6.1 | 7.15 |
| studious | 6.8 | 4.45 | 6.55 |
| tolerant | 7 | 3.45 | 6.85 |
| respectful | 6.95 | 3.9 | 6.75 |
| loyal | 6.9 | 4 | 6.8 |
| happy | 7.9 | 5.5 | 7.2 |
| earnest | 6.45 | 3.95 | 5.9 |
| pleasant | 7.65 | 5 | 6.95 |
| appreciative | 7.45 | 5 | 7.2 |
| generous | 7 | 3.7 | 6.55 |
| mature | 6 | 3.55 | 6.45 |
| tidy | 7.1 | 3.85 | 7.1 |
| fearless | 6.55 | 4.7 | 6.1 |
| unselfish | 7 | 4.2 | 6.75 |
| interesting | 7.35 | 5.7 | 7 |
| tactful | 6.95 | 4.4 | 5.5 |
| sharp-witted | 6.1 | 4.5 | 5.4 |
| decisive | 6.4 | 4.3 | 5.6 |
| righteous | 6.6 | 4 | 6.75 |
| decent | 6.3 | 3.7 | 6.05 |
| vivacious | 7.4 | 6.1 | 6.65 |
| gentle | 6.75 | 2.95 | 6.95 |
| warm | 6.95 | 3.75 | 6.95 |
| obliging | 7.3 | 5.05 | 6.8 |
| enthusiastic | 7.4 | 6.3 | 6.7 |
| self-reliant | 6.1 | 4.15 | 6.6 |
| rational | 6.3 | 3.25 | 6.85 |
| direct | 6.5 | 4.3 | 6.5 |
| sincere | 7.25 | 4.4 | 7.1 |
| active | 7.2 | 5.45 | 6.85 |
| meticulous | 6.15 | 4.25 | 6.4 |
| bright | 7.1 | 4.45 | 6.3 |
| smart | 7 | 4.15 | 6.8 |
| intelligent | 6.95 | 4.45 | 6.05 |
| capable | 6.95 | 4.1 | 6.2 |
| self-confident | 7.3 | 4.85 | 6.5 |
| self-contented | 6.7 | 5.3 | 6.3 |
| thrifty | 6.1 | 4.05 | 6.55 |
| honest | 7 | 3.4 | 7.1 |
| humble | 6.85 | 3.15 | 6.8 |
| modest | 7.25 | 3.55 | 7.25 |
| cautious | 6.2 | 3.6 | 6.95 |
| enterprising | 6.9 | 5.45 | 6.6 |
| easygoing | 6.7 | 3.7 | 6.75 |
| amusing | 7.1 | 5.3 | 6.2 |
| resentful | 2.35 | 5.2 | 6.85 |
| domineering | 2.5 | 5.05 | 5.15 |
| insolent | 3.2 | 4.6 | 5.55 |
| offensive | 2.7 | 5.85 | 6.25 |
| cold | 3.2 | 2.75 | 6.75 |
| messy | 3.75 | 4.35 | 6.25 |
| unkind | 2.85 | 4.55 | 5.6 |
| snobbish | 3.15 | 4.05 | 5.55 |
| mean | 2.85 | 3.65 | 6.1 |
| absent-minded | 3.95 | 3.8 | 6.95 |
| obstinate | 3.5 | 5.35 | 6.8 |
| discourteous | 2.55 | 5.4 | 5.95 |
| negligent | 2.25 | 5.65 | 5.65 |
| envious | 2.75 | 5.2 | 6.5 |
| antisocial | 3.05 | 2.9 | 6.3 |
| stingy | 2.75 | 4.15 | 6.25 |
| childish | 3.85 | 4.85 | 6.3 |
| weak | 2.6 | 3.9 | 6.05 |
| distrustful | 3.5 | 4.25 | 6.35 |
| hot-tempered | 3.85 | 6.5 | 6.75 |
| malicious | 2.4 | 6 | 5.35 |
| pessimistic | 2.15 | 3.75 | 6.05 |
| unwise | 3 | 4.25 | 6 |
| foolish | 3.25 | 4.15 | 6.1 |
| listless | 3.95 | 3.3 | 6.75 |
| cowardly | 2.65 | 3.9 | 6.25 |
| stubborn | 3.05 | 4.55 | 5.15 |
| complaining | 2.9 | 5.1 | 6.45 |
| fault-finding | 2.9 | 5.15 | 6.3 |
| helpless | 2.45 | 6.35 | 5.7 |
| unreasonable | 2.4 | 4.85 | 6.8 |
| ungracious | 3.4 | 3.85 | 6.25 |
| irritable | 2.7 | 6.65 | 6.05 |
| cruel | 2.6 | 6.65 | 6.65 |
| shallow | 2.35 | 4 | 5.9 |
| heartless | 2.65 | 6.15 | 6.2 |
| sly | 3.05 | 5.6 | 5.8 |
| deceitful | 3.2 | 5.1 | 5.9 |
| dominating | 2.75 | 5.4 | 5.3 |
| hostile | 2.45 | 5.55 | 5.75 |
| neglectful | 3.25 | 4.9 | 6.3 |
| thoughtless | 3.1 | 5.1 | 6.5 |
| impolite | 3 | 4.85 | 6.2 |
| nervous | 3.15 | 6.9 | 6.15 |
| superficial | 3.2 | 3.85 | 6.1 |
| timid | 3 | 4.8 | 5.7 |
| conceited | 3.5 | 5.35 | 5.75 |
| boastful | 3.5 | 4.8 | 5.25 |
| selfish | 2.8 | 5.1 | 6.55 |
| self-conceited | 2.9 | 4.65 | 5.5 |
| sloppy | 3.65 | 5.05 | 6 |
| insincere | 2.7 | 5.15 | 5.85 |
| untruthful | 3 | 4.9 | 5.35 |
| greedy | 2.95 | 5 | 5.65 |
| scornful | 3.2 | 4.5 | 5 |
| untidy | 3.15 | 4.3 | 5.45 |
| gloomy | 3.15 | 3.55 | 5.4 |
| deceptive | 3.05 | 5.3 | 5.95 |
| hot-headed | 3.5 | 5 | 5.6 |
| troublesome | 2.7 | 5.25 | 6.2 |
| positive | 7.05 | 3.7 | 6.6 |
